# SOD2 in Skeletal Muscle: New Insights from an Inducible Deletion Model

**DOI:** 10.1101/2021.05.20.443346

**Authors:** Aowen Zhuang, Christine Yang, Yingying Liu, Yanie Tan, Simon T. Bond, Shannen Walker, Tim Sikora, Arpeeta Sharma, Judy B. de Haan, Peter J. Meikle, Melinda T. Coughlan, Anna C. Calkin, Brian G. Drew

## Abstract

Metabolic conditions such as obesity, insulin resistance and glucose intolerance are frequently associated with impairments in skeletal muscle function and metabolism. This is often linked to dysregulation of homeostatic pathways including an increase in reactive oxygen species (ROS) and oxidative stress. One of the main sites of ROS production is the mitochondria, where the flux of substrates through the electron transport chain (ETC) can result in the generation of oxygen free radicals. Fortunately, several mechanisms exist to buffer bursts of intracellular ROS and peroxide production, including the enzymes Catalase, Glutathione Peroxidase and Superoxide Dismutase (SOD). Of the latter there are two intracellular isoforms; SOD1 which is mostly cytoplasmic, and SOD2 which is found exclusively in the mitochondria. Developmental and chronic loss of these enzymes has been linked to disease in several studies, however the temporal effects of these disturbances remain largely unexplored. Here, we induced a post-developmental (8-week old mice) deletion of SOD2 in skeletal muscle (SOD2-iMKO) and demonstrate that 16 weeks of SOD2 deletion leads to no major impairment in whole body metabolism, despite these mice displaying alterations in aspects of mitochondrial abundance and voluntary ambulatory movement. Furthermore, we demonstrated that SOD2 deletion impacts on specific aspects of muscle lipid metabolism, including the abundance of phospholipids and phosphatidic acid (PA), the latter being a key intermediate in several cellular signaling pathways. Thus, our findings suggest that post-developmental deletion of SOD2 induces a more subtle phenotype than previous embryonic models have shown, allowing us to highlight a previously unrecognized link between SOD2, mitochondrial function and bioactive lipid species including PA.

## Introduction

Insulin resistance, glucose intolerance and type 2 diabetes are strongly influenced by a combination of genetics and lifestyle, which is further compounded by the process of ageing^1^. Many pathways have been identified as contributing determinants of these diseases, however one of the most consistent, reproducible features amongst them all is mitochondrial dysfunction^2^. This suggests a direct association between mitochondrial function in peripheral tissues and the progression of metabolic disease, however which of these conditions is the primary driving factor remains to be determined. In an effort to tease out this relationship, studies have suggested that early insults that directly impact on mitochondria, such as mutations in key mitochondrial genes, promote mitochondrial dysfunction and induce type 2 diabetes^3^.

When mitochondria are dysfunctional they are often less efficient at generating ATP through oxidative phosphorylation, and thus several toxic by-products are generated including excessive amounts reactive oxygen species (ROS). Whilst ROS in normal concentrations are important cellular signaling molecules, excessive and chronic production of ROS can lead to deleterious effects including oxidation of proteins and metabolites, mutation of mitochondrial DNA, inhibition of glycolysis and the promotion of advanced glycation end products^4^. Fortunately, in a healthy cell several mechanisms exist to buffer these transient bursts of ROS, including the activity of intracellular enzymes such as Catalase and Glutathione Peroxidase (Gpx1), and two isoforms of Superoxide Dismutase (SOD); SOD1 and SOD2^5^.

The ROS buffering capacity of a cell can, however, be significantly reduced by several mechanisms including mutations in genes important for ROS production or scavenging, or perturbations to energy substrate metabolism. For example, when a cell is exposed to high levels of glucose, this forces excess substrate through the electron transport chain (ETC), placing a greater demand on resources^4^. Inevitably, the ETC becomes overloaded and unless there is an outlet for the increased electrons produced along the pathway such as uncoupling proteins, they are instead shunted onto molecular oxygen which chemically reduces it to produce superoxides, or ROS. This increase in ROS further reduces mitochondrial capacity and thus sets in place a vicious cycle whereby increasing glucose concentrations continues to elevate levels of ROS, and vice versa^5^.

Depending on the tissue in which mitochondrial dysfunction and chronic ROS accumulation occurs, it can manifest in different phenotypes. Chronic increases in ROS promotes cardiomyopathy, chronic kidney disease and skeletal muscle atrophy and perturbed energy metabolism^6^. All of these phenotypes are common in the setting of glucose intolerance and diabetes, suggesting that prevention or treatment of pathways that lead to elevated ROS, is of potential therapeutic interest.

Many groups have been interested in the notion that reducing excessive ROS is a means to preventing disease. Indeed, several pharmaceutical companies have made significant investments into developing modified antioxidant molecules such as MitoQ, MTP-131 and SkQ1 that have demonstrated various levels of efficacy in human trials of sarcopenia, cardiac disease and fatty liver disease^7^. Moreover, preclinical genetic models have elegantly demonstrated that global overexpression of antioxidant enzymes such as catalase and SOD, which aid in mopping up excess ROS, have efficacy in preventing endothelial dysfunction, atherosclerosis, fatty liver, heart disease and glucose intolerance^8,9^. On the contrary, global genetic deletion of these enzymes induce pathological phenotypes. For example, homozygous deletion of SOD2 (Mn-SOD), a variant of SOD found exclusively in the mitochondria, led to neonatal lethality^10^. Furthermore, although viable, heterozygous SOD2 mutants present with many pathologies consistent with mitochondrial dysfunction including poor tolerance to exercise, cardiomyopathy, cardiovascular disease and glucose intolerance^9,11–15^. Thus, manipulating cellular ROS either through pharmacological or genetic means appears to impact on disease outcomes. A potential downside with the majority of studies thus far, has been the inability to tease out what the tissue specific effects of ROS damage are, which might aid in improving antioxidant targeting for therapeutic benefit.

One enzyme studied for its tissue specific effects is SOD2, with liver^16^, brain^17^, adipose^18^, heart^19^, smooth muscle^20^ and skeletal muscle^14,15^ specific models all having been generated and phenotyped. Interestingly, the skeletal muscle specific studies have demonstrated striking defects in exercise capacity, muscle strength and mitochondrial activity, similar to what has been observed in whole body models^14,15^. An important point to note is that these prior models deleted SOD2 during early development, raising questions as to whether the observed phenotypes were due to increased mitochondrial ROS in fully matured muscle *per se*, or whether in fact the loss of SOD2 and a subsequent increase in ROS in the developmental stages, impacted skeletal muscle development directly. Thus, studies which aim to understand the effect of SOD2 deletion in skeletal muscle post-development, stand to overcome this limitation and shed new light on our understanding of this pathway.

In this study we investigated the molecular and metabolic effects of SOD2 deletion from the musculature in post-developmental male mice from 8 weeks of age.

## Results

### Generation and validation of post-developmental, skeletal muscle specific SOD2-knock out mice

To generate mice with deletion of SOD2 in skeletal muscle post-development, we crossed SOD2 floxed (SOD2 fl/fl) mice with ACTA1-creERT2 (mCre) mice. The resulting SOD2 fl/fl and fl/fl-creERT2 (fl/fl-mCre) male mice were subsequently administered Tamoxifen (TAM; 80mg/kg) in sunflower oil treated by gavage, or with sunflower oil alone for 3 consecutive days at approximately 8 weeks of age to activate cre-recombinase. Mice were then fed a high fat diet for the subsequent 12 weeks to metabolically stress the animals. A separate cohort of fl/fl and fl/fl-mCre mice were treated with tamoxifen, and fed a normal chow diet for 12 weeks (no vehicle control mice for chow study). High fat diet (HFD) mice underwent a comprehensive phenotyping regimen over the 12-week period before collecting tissues at the end of the study following a 6 hour fast. Blood and tissues were also collected from chow fed mice at study end.

Using qPCR analysis (see **Table 1** for qPCR primers sets) on muscle acquired from HFD fed mice, we demonstrated that cre-recombinase expression was specific to skeletal muscle, with no expression detected in liver or white adipose tissue (WAT) (**Figure 1A**). Moreover, SOD2 mRNA expression was almost completely ablated in skeletal muscle, whereas no change was observed in liver or WAT (**Figure 1B**). To investigate the effect of cre-lox activity in different muscle tissues, we investigated cre-recombinase and SOD2 mRNA expression in a mixed fibre-type muscle (*Tibialis anterior* – TA), a red fibre-type muscle (*soleus*) and a white fibre-type muscle (*Extensor digitorum longus* – EDL). These results demonstrate that cre-expression appears to be lower in the *Soleus* red muscle fibre type compared to TA and EDL (**Figure 1C**), however the deletion of SOD2 was equivalent between all three muscle types (**Figure 1D**), validating SOD2-deletion in all muscle types. To confirm that ablation of SOD2 mRNA resulted in deletion of the SOD2 protein, we performed western blots on TA muscles from both chow and high-fat diet (HFD) fed mice. This demonstrated almost complete ablation of SOD2 in fl/fl-mCre (KO) muscle compared to fl/fl mice (WT) (**Figure 1E**).

**Table 1.**
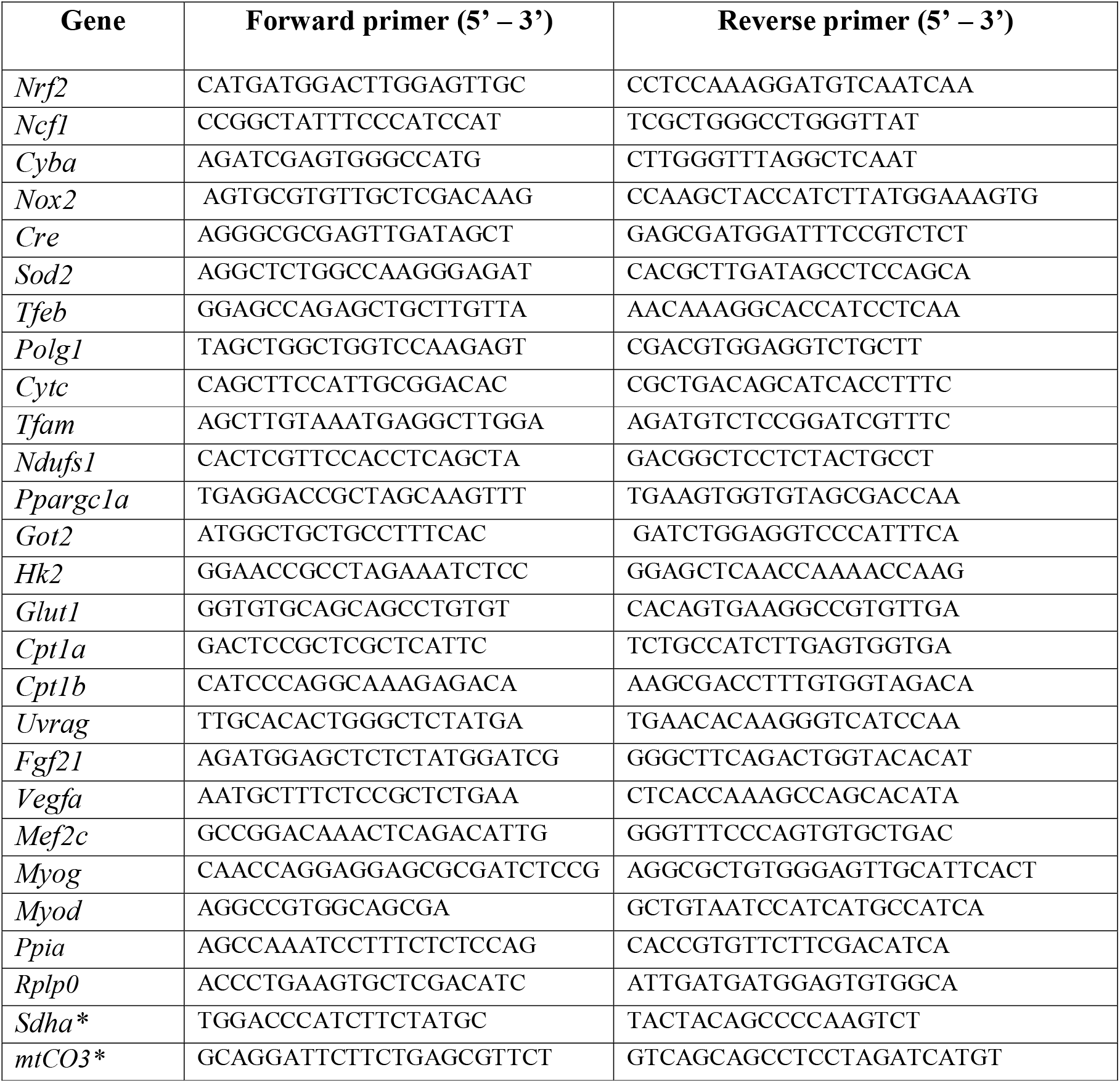
Forward and reverse primer sets for detection of the designated gene using qPCR. (m = mouse). * indicates primer sets used on DNA, to determine the mtDNA/tDNA ratio.

**Figure 1:**
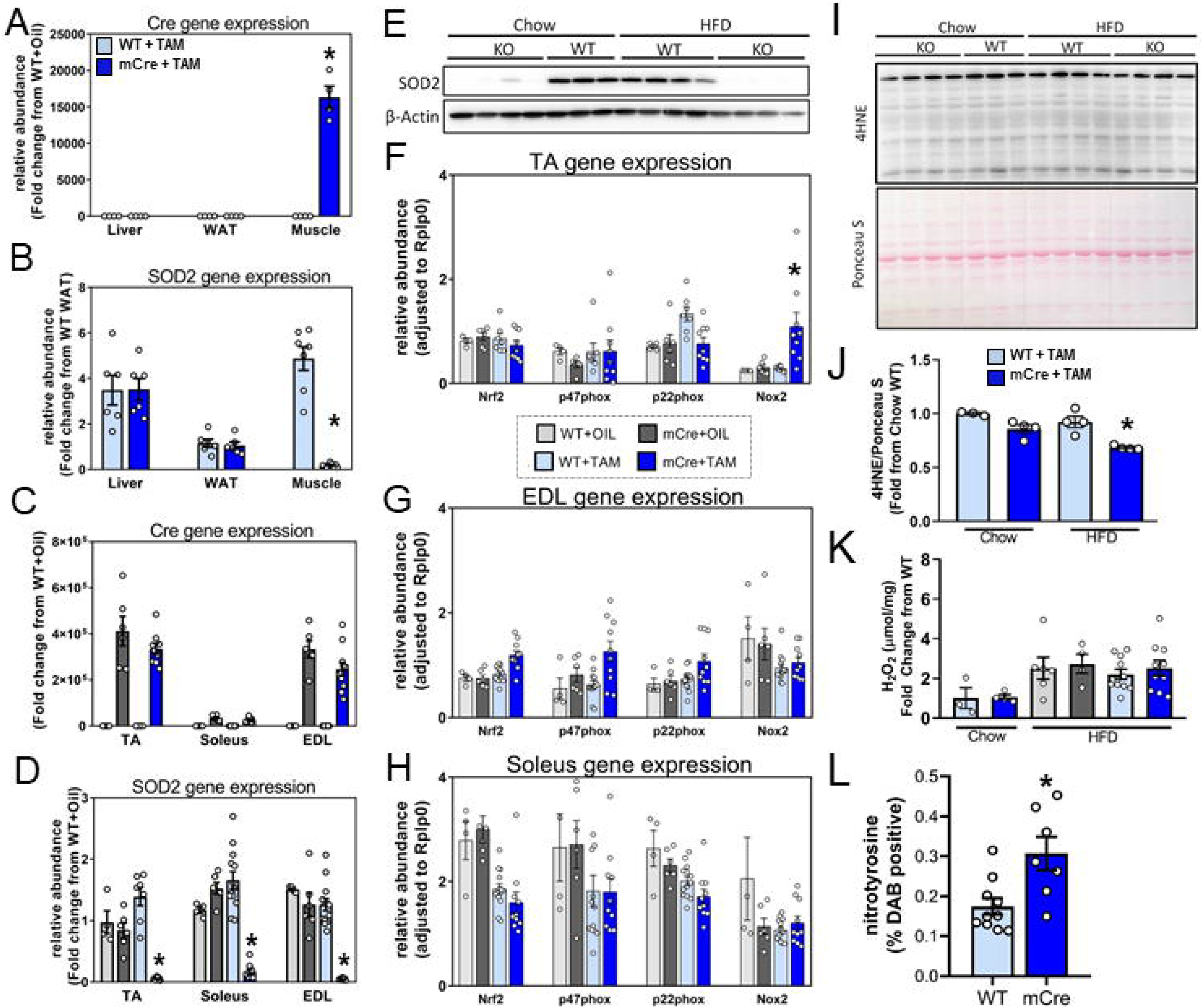
Generation and characterisation of inducible, skeletal muscle specific SOD2 knock out mice. mRNA expression of **(A)** cre recombinase and **(B)** SOD2 in liver, white adipose tissue (WAT) and TA muscle of SOD2 WT and mCre mice (n=4/group); **(C)** Cre recombinase and **(D)** SOD2 mRNA expression across three different muscle types (TA, Soleus and EDL) of WT and mCre positive mice (n=6-11/group); **(E)** Western blots for SOD2 and β-actin in TA muscle lysates from WT plus tamoxifen (WT) and mCre plus tamoxifen (KO) mice fed either a chow or high fat diet (HFD; n=3-4/group). mRNA expression of redox genes from **(F)** TA, **(G)** EDL and **(H)** soleus muscles from the four HFD fed cohorts (n=4-11/group); **(I)** Western blot and **(J)** quantitation of 4-hydroxynonenal (4HNE) conjugated proteins from TA of WT plus tamoxifen (WT) and mCre plus tamoxifen (KO) mice fed either a chow or HFD (n=3-4/group); **(K)** Skeletal muscle peroxide abundance as determined by Amplex Red assay in TA muscles from WT and KO chow fed mice and WT+OIL, mCre+OIL, WT+TAM, mCre+TAM cohorts fed a chow or HFD (n=3-10/group); **(L)** Quantification of Nitrotyrosine staining as determined by immunohistochemistry performed on sections of TA muscles from WT+TAM and mCre+TAM fed a HFD (n=7-10/group) All data are presented as mean ± SEM. Data with >2 groups were compared by ANOVA with Fishers LSD post-hoc testing, samples with 2 groups were analysed using a Mann-Whitney non-parametric t-test where * denotes a p-value<0.05. WAT = white adipose tissue, TA = *Tibialis anterior*, EDL = *Extensor digitorum longus*, 4HNE = 4-hydroxynonenal, DAB = 3,3′-Diaminobenzidine; HFD = high fat diet.

Upon demonstrating that SOD2 mRNA and protein was specifically deleted in the skeletal muscles of fl/fl-mCre mice, we next sought to investigate the effect of SOD2 deletion on readouts of redox regulation in skeletal muscle. This was performed on three different muscles (TA, EDL and Soleus) from HFD fed animals, across all 4 groups of mice; WT+OIL, mCre+OIL, WT+TAM and mCre+TAM. We initially performed gene expression analysis using qPCR for known enzymes that are involved in redox regulation including *Nrf2*, *p47Phox*, *p22Phox* and *Nox2* (**Figures 1F–H)**. These genes were largely unaffected by the deletion of SOD2 (dark blue bars), except for *Nox2* which was significantly increased in TA of KO mice (**Figure 1H, dark blue bar**), whilst *Nrf2* and *p47phox* showed a trend to be increased in the EDL of KO mice (**Figure 1G, dark blue bar**). No genes were altered by the deletion of SOD2 in the soleus, although it did appear that Tamoxifen treatment alone induced a general reduction in each of the genes assayed (**Figure 1H**, blue bars).

Next we investigated the effect of SOD2 deletion on the abundance of reactive oxygen species (ROS), and downstream products known to be altered by oxidative stress. **Figure 1I** shows the abundance of 4-hydroxynonenal (4HNE) protein adducts as determined by Western blot, which are often generated as a result of increased free radicals. Skeletal muscle deletion of SOD2 resulted in a reduction of 4HNE staining in TA muscles from both chow and HFD fed animals, which was significantly (p=0.004) different in the HFD fed animals (**Figure 1J)**, indicating a reduced protein peroxidation in the absence of SOD2. We also performed a direct measure of peroxides (e.g. H_2_O_2_) in the TA muscles using an Amplex Red Assay, which demonstrated an increase in the abundance of peroxides in the skeletal muscle of HFD fed animals compared to chow fed animals. However, SOD2 KO did not have any impact on peroxide abundance (**Figure 1K**), which is somewhat surprising given that the role of SOD2 is to convert mitochondrial superoxides and thus one might have expected a decrease in peroxide concentration in the absence of SOD2. Given there was no change in the level of peroxide in SOD2 KO compared to WT HFD fed mice, this would suggest that the majority of peroxide generated in this setting are not derived from the mitochondria, and is likely impacted by alterations in other enzymes including SOD1. Finally, we measured the amount of nitrotyrosine products generated in TA muscles of WT and SOD2-KO (mCre) HFD fed mice. This demonstrated a ∼2-fold increase in positive staining for nitrotyrosine in muscle sections of SOD2 KO muscle (p<0.05; **Figure 1L**), suggestive of increased superoxide levels in SOD2 KO that lead to increased protein nitration via peroxynitrite intermediates. Together, these data confirm that post-developmental deletion of SOD2 leads to alterations in redox readouts in skeletal muscle, with nitrotyrosine noticeably increased in KO muscles.

### Inducible, muscle specific knock out of SOD2 has no effect on body mass or organ weights

Upon demonstrating successful generation of muscle specific SOD2 KO mice with mild alterations to redox homeostasis, we sought to investigate if feeding these mice a HFD led to genotype specific alterations in body mass and tissue weights. Weekly analysis of body weight revealed that KO mice (mCre+TAM) had a similar weight gain over the 12 week study compared to the three control models (WT+OIL, mCre+OIL, WT+TAM) (**Figure 2A**). This was confirmed by EchoMRI that demonstrated no difference in lean mass (**Figure 2B**) or fat mass (**Figure 2C**) between all groups at any of the timepoints measured; however, it did demonstrate marked increase in fat mass in all models in response to HFD, as expected. With regard to organ weights at study end, we demonstrated no difference in EDL (**Figure 2D**), TA (**Figure 2E**), liver (**Figure 2F**) or gonadal white adipose tissue (gWAT) (**Figure 2G**) weight across the four cohorts. Overall, these data demonstrated that inducible, muscle specific SOD2 KO does not alter body mass or tissue weights in mice fed a HFD for 12 weeks.

**Figure 2:**
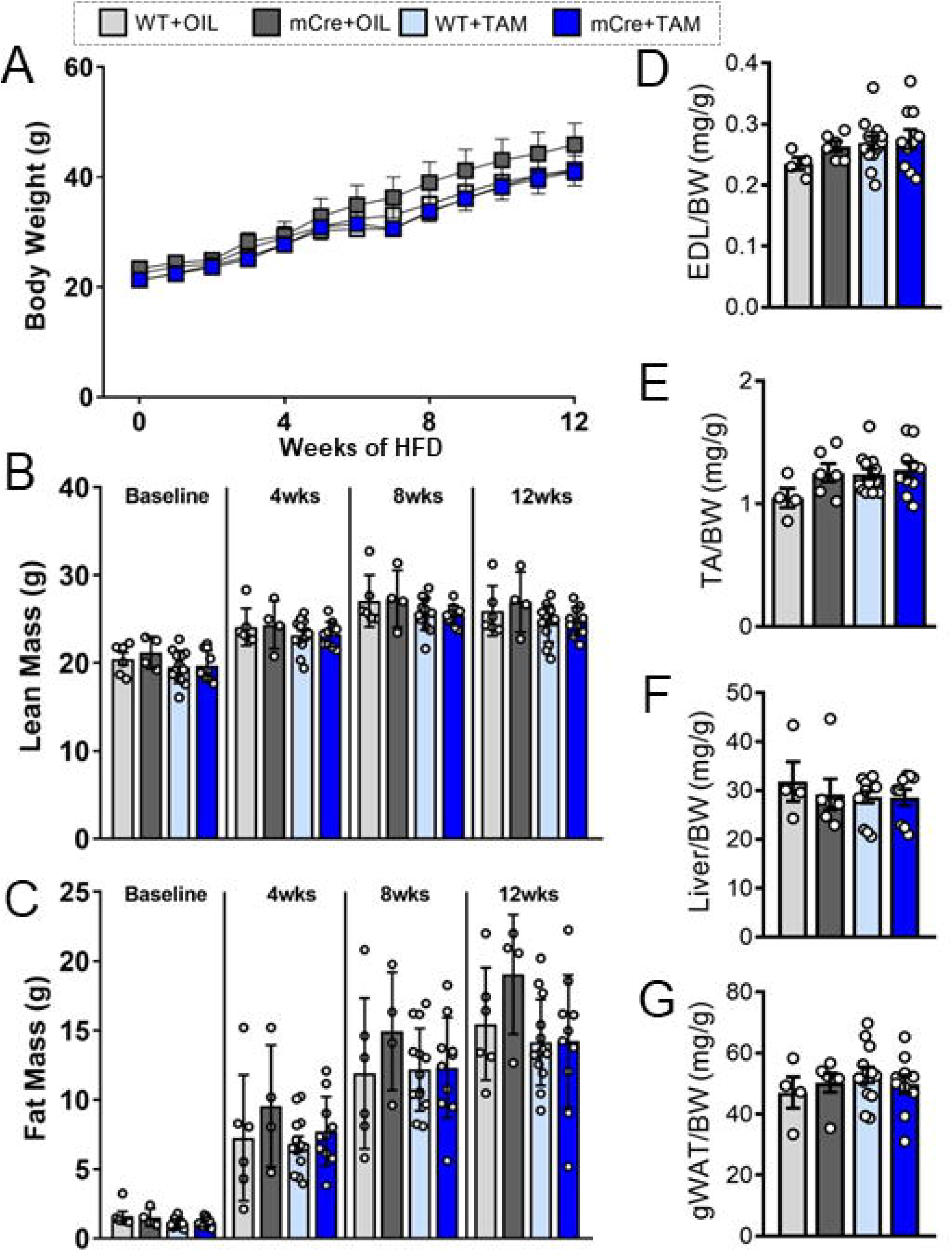
General features and tissue weights of inducible, skeletal muscle specific KO mice. **(A)** Weekly total body weight of the four cohorts fed a HFD for 12 weeks (n=4-11/group); **(B)** Lean mass and **(C)** fat mass as determined by EchoMRI in the four HFD fed cohorts at baseline, 4 weeks, 8 weeks and 12 weeks post-HFD (n=4-11/group); Tissue weight for **(D)** EDL, **(E)** TA, **(F)** liver and **(G)** gonadal WAT (gWAT) normalized to body weight (BW) at study end (n=4-11/group). Data are presented as mean ± SEM. Data in (A-C) were analysed using repeated measures two-way ANOVA. All other data were compared by ANOVA with Fishers LSD post-hoc testing. g = grams, mg = milligrams, wks = weeks, HFD = high fat diet, TA = *Tibialis anterior*, EDL = *Extensor digitorum longus*, gWAT = gonadal white adipose tissue, BW = body weight.

### Inducible, muscle specific knock out of SOD2 alters ambulatory movement behaviour, but does not affect glucose tolerance

After demonstrating that inducible deletion of SOD2 in skeletal muscle had no effect on total body mass or organ weights, we sought to determine if subclinical changes in metabolism and movement behavior were apparent in these mice. To study these outputs, we placed the WT+TAM and mCre+TAM cohorts in a Promethion High-Definition Multiplexed Respirometry System both prior to commencement of diet (baseline) and after 10 weeks of HFD feeding (End HFD). The Promethion system performs repeated measures of whole body respiration, activity behavior and daily movement of individual mice across the entire 24-hour analysis period. These studies demonstrated that the respiratory exchange ratio (RER) between the two genotypes was not different at any particular period over the 24 hours (**Figure 3A**). There was also no difference in RER between genotypes at baseline or after HFD (**Figures 3B&3C**), or in energy expenditure (EE) (**Figures 3D&3E**). However, as expected there was a drop in the RER observed between baseline and HFD, particularly during the night period (∼0.85 to 0.78 at light, ∼0.93 to 0.6 at dark, for NC vs HFD respectively), consistent with the animals using more fat for energy production. HFD feeding also coincided with an increased EE after HFD feeding compared to baseline, which was particularly noticeable during the light cycle, likely reflecting the high-energy content of the diet they were consuming.

**Figure 3:**
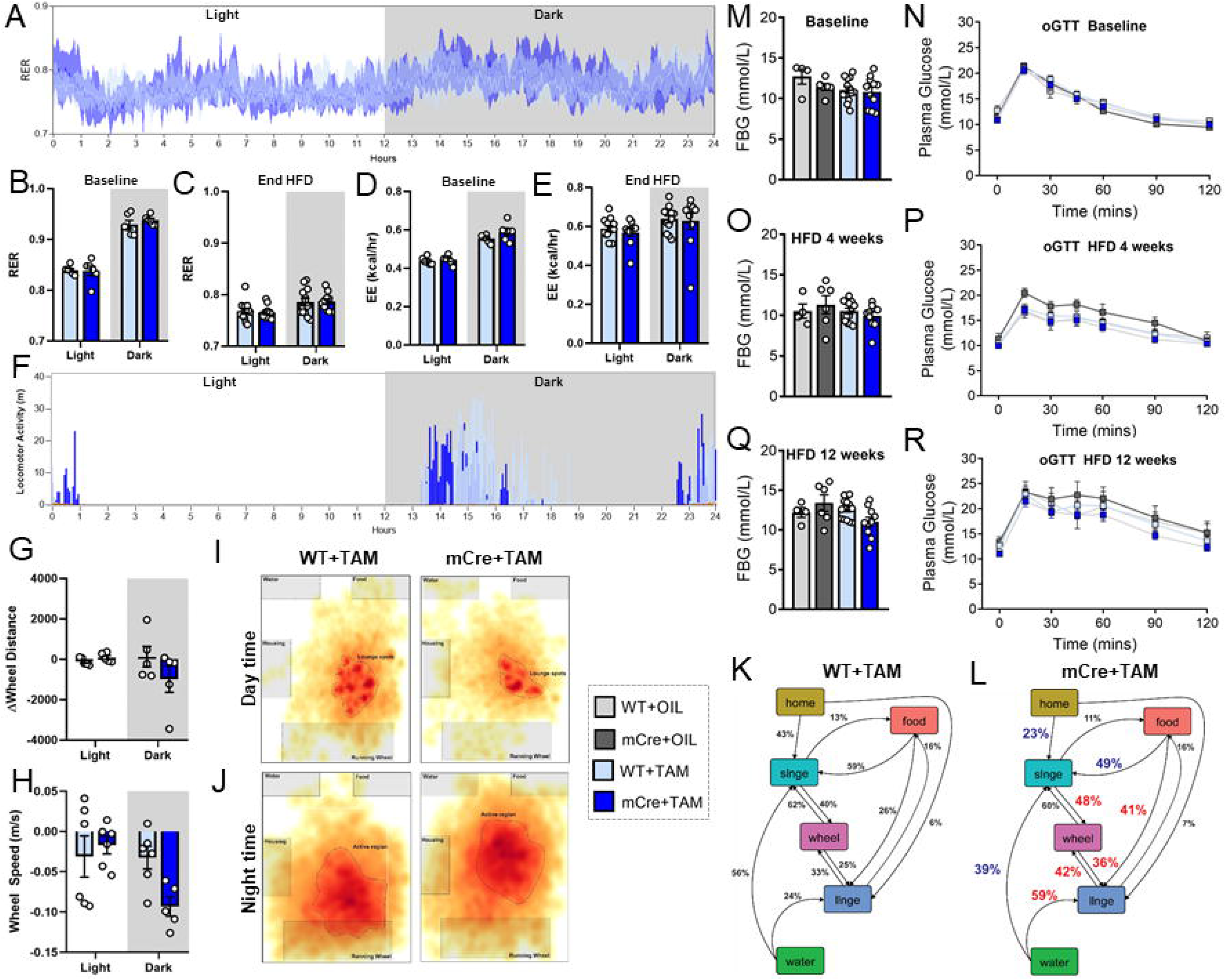
Measurements of whole body metabolism and glucose tolerance in inducible, muscle specific SOD2 KO mice. **(A)** Respiratory exchange ratio (RER) trace from Promethion analysis over the entire 24 hour assessment period, **(B)** average RER and **(C)** average energy expenditure (EE) for WT+TAM (light blue) and mCRE+TAM (dark blue) mice in the light and dark periods; **(D)** Average RER and **(E)** average energy expenditure (EE) for both genotypes in the light and dark periods prior to study end; **(F)** Total locomoter activity (pedal and wheel movement) over the 24 hour assessment period at the end of the HFD feeding study; Change in **(G)** average wheel distance and **(H)** average wheel speed from baseline to study end during the light and dark cycles; Locality mapping of mice based on 24 hours of movement data during the **(I)** light cycle (day time) and **(J)** dark cycle (night time). Map colours indicate increasing time spent in that given location (red being most, yellow being least). Intense red areas indicate “lounge spots”, likely representing regions where mice spent time resting for longer periods. Grey boxes designate each fixed cage component (i.e. wheel, food hopper, water sipper and running wheel). Probability mapping for **(K)** WT+TAM mice and **(L)** mCRE+TAM mice estimates predicted behavior (percentage) following a certain activity. Red numbers indicate an increase, whilst blue numbers indicate a decrease in that behavior in KO mice relative to WT mice, *slnge* = short lounge, *llnge* = long lounge. Fasting blood glucose (FBG) measurement and oral glucose tolerance tests (oGTT) in mice at **(M, N)** baseline, **(O, P)** 4 weeks post-HFD and **(Q, R)** 12 weeks post-HFD respectively. Data are presented as mean ± SEM (B-E, G, H, M-R). Data in (N, P and R) were compared using repeated measures ANOVA. All other data were compared by ANOVA with Fishers LSD post-hoc testing where * indicates a p-value<0.05. RER = respiratory exchange ratio, EE = energy expenditure, HFD = high fat diet.

Further to respiration and energy expenditure measures, the Promethion system is able to provide insights into animal behavior and ambulatory movement. We observed a noticeable shift in the times and pattern of active movement in the KO mice compared to WT mice (**Figure 3F**). In this panel, it was observed that KO animals tended to do more intense movement early on in the night cycle, when they first wake, compared to WT mice. Following this bout of movement, they tended not to move again substantially until the end of the night cycle, which then continued well into the early part of the day cycle. These altered behaviors encouraged us to investigate this finding further, and thus we studied their locomotion and exercise behaviours. To do this we compared the distance and speed at which they travelled on the running wheel at the start of the study (baseline), with that at the end of the study (end of HFD). This was performed for both genotypes and separated into light and dark activity cycles. Thus, if the distance and speed the mice covered was the same at the beginning and the end of the study, the “delta” would be zero. We demonstrated that there was no difference in the distance both genotypes ran at the start and the end of study during the day time cycle (**Figure 3G)**, nor was there a difference in WT mice in the night time cycle. However, there was a strong trend for a reduced distance covered by KO mice in wheel distance in the night time cycle, as indicated by the negative delta. With regards to wheel speed, we demonstrated that both genotypes were unable to maintain the same speed at the end of the study compared to their baseline speeds (perhaps due to the HFD), however there was a noticeable and robust decline in wheel speed in the KO mice in the nighttime cycle compared to WT mice (**Figure 3H)**. These data collectively demonstrate that KO mice have a decline in the capacity to travel the same distance, and at the same speed as WT mice.

Another measure we can obtain from Promethion is positional probability mapping, which uses data collected over the entire analysis period to estimate cumulative animal movement behaviours. This analysis demonstrated that KO mice spend more time “lounging” in specific regions of the cage for long periods of time than WT mice (**Figure 3I and 3J**). This can be observed both in the day time (**Figure 3I)**, where less lounge spots for the KO mice indicate longer time spent in the one spot, and at night time (**Figure 3J)** where the KO mice spend far less time near the wheel and closer to lounge spots. In an attempt to quantify these observations, we used behavioral transition analysis to infer the percent of time the different genotypes spent doing each activity (**Figure 3K&3L)**. The propensity for KO mice to take a long lounge (llnge) appeared to be more apparent after they had either eaten food, drank water or undertaken exercise (KO vs WT: 41% vs 26%, 36% vs 25%, 59% vs 24% respectively), suggestive of fatigue (increased percentage shown in red). Indeed, WT mice were more likely to take a short lounge (slnge) after eating or drinking, supporting the notion that KO mice were more lethargic and less willing to stay active after these activities.

To determine whether these alterations in ambulatory movement and animal behavior impacted on metabolism of energy substrates in these mice, we assessed their whole body glucose regulation. At baseline there were no differences in either basal fasting glucose (**Figure 3M**), or in their tolerance to a standardized bolus of glucose (**Figure 3N**). Indeed, this remained similar throughout the HFD feeding regimen where, despite glucose tolerance deteriorating as expected over the HFD feeding period, it remained similar between the KOs and all three control models at both 4 weeks and 12 weeks post diet (**Figures 3O–3R**). Thus, although the movement and behavior of the KO mice was altered, this did not appear to impact on glucose tolerance in these animals, even following a 12-week HFD challenge.

### Inducible, muscle specific knock out of SOD2 induces changes in skeletal muscle mitochondrial composition

Given that SOD2 is localized to the mitochondria, and that we observed phenotypes in KO mice that were reminiscent of lethargy that is often observed in models with mitochondrial deficits, we performed experiments to test aspects of mitochondrial activity. We initially performed analysis on the abundance of mtDNA in the TA muscle of all four cohorts from the HFD study (**Figure 4A**). These data demonstrated that mtDNA abundance was not different between KO mice and the various control cohorts. Next, we performed Western blot analysis on the TA muscles from WT+TAM and mCre+TAM animals fed both chow and HFD to investigate the abundance of representative proteins from each of the five complexes of the electron transport chain (ETC) (**Figure 4B**). These data demonstrated that SOD2 KO mice had a substantial reduction in the abundance of both Complex I and Complex II proteins of the ETC (**Figure 4C**). Moreover, although not significant, there was a notable trend for a reduction in all other complexes of the ETC. Given the robust reductions in ETC complex I and II, we performed gene expression analysis on the three different muscle types (TA, EDL, Soleus) from all four cohorts of mice that were fed a HFD, to determine if this phenotype might be driven by transcriptional changes in mitochondrial genes (**Figures 4D–4F**). These data demonstrated that although some genes were modestly changed in KO TA muscles (e.g. *Ppargc1a* and *Ndufs1*), overall the expression of key mitochondrial genes was not altered by deletion of SOD2. These findings indicated that key components of the mitochondrial ETC were reduced in SOD2 KO skeletal muscles, but this was not due to changes in mtDNA abundance or changes in gene transcription, suggesting a post-translational effect on ETC complex abundance in this model potentially due to altered redox homeostasis.

**Figure 4:**
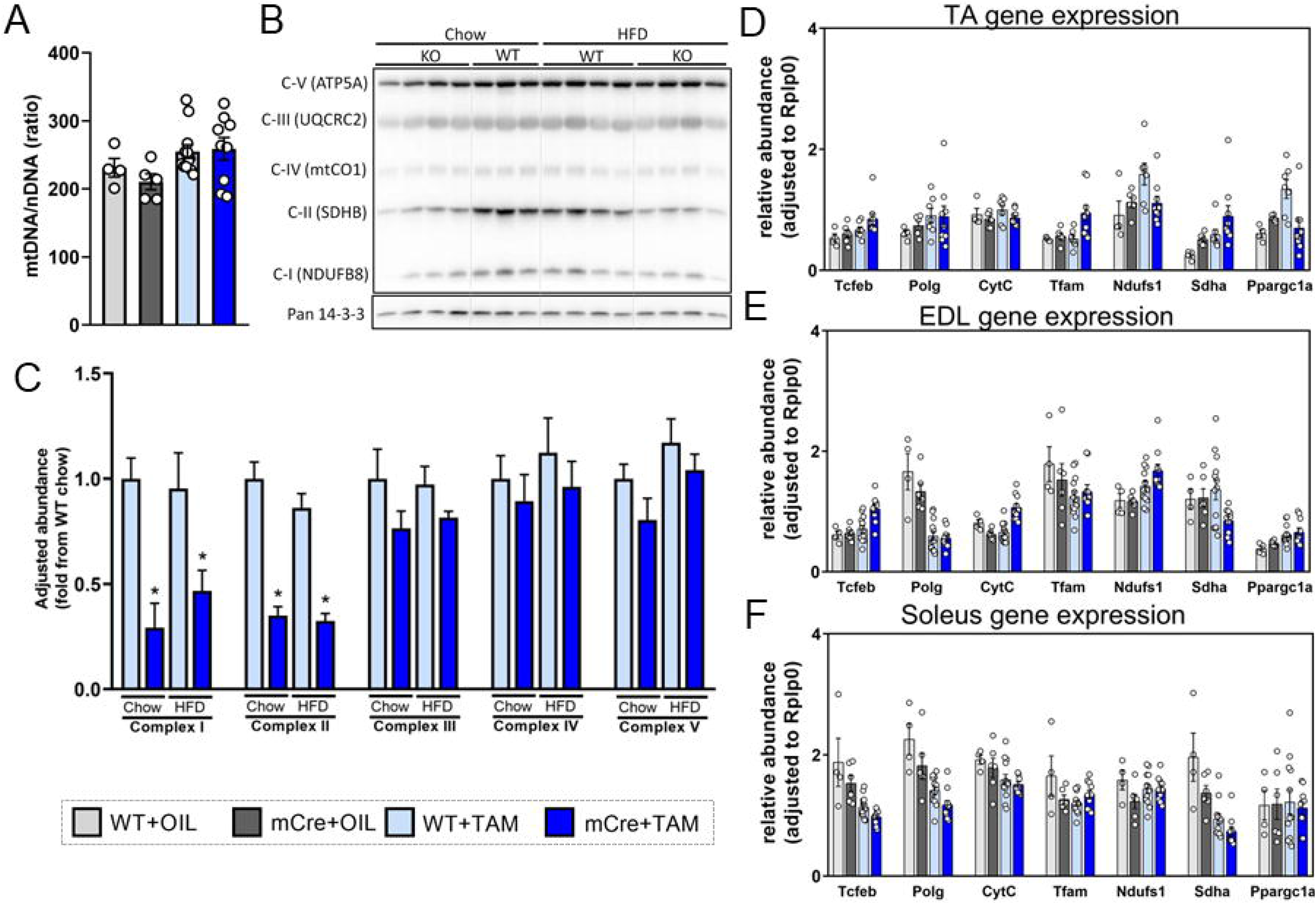
Analysis of mitochondrial abundance and gene expression in muscles from inducible, skeletal muscle specific SOD2 KO mice. **(A)** Mitochondrial DNA (mtDNA) to nuclear DNA (nDNA) ratio in TA muscles of the four HFD fed cohorts; **(B)** Western blot on protein from TA muscles isolated from WT+TAM (WT) and mCre+TAM (KO) mice fed a chow diet or high fat diet (HFD) for proteins from the 5 complexes (CI-CV) of the mitochondrial electron transport chain (ETC) and of pan 14-3-3 (loading control). **(C)** Quantification of five proteins from complexes of the ETC from panel B, normalized to the the loading control (14-3-3); qPCR analysis for genes involved in skeletal muscle mitochondrial function from **(D)** TA **(E)** EDL and **(F)** Soleus muscles from all four cohorts of HFD fed mice. All data are presented as mean ± SEM. All data were compared by ANOVA with Fishers LSD post-hoc testing where * indicates a p-value<0.05. TA = *Tibialis anterior*, EDL = *Extensor digitorum longus*.

### Inducible, muscle specific knock out of SOD2 alters pathways involved in energy substrate utilisation

Given the subtle alterations in mitochondrial readouts, we speculated whether SOD2 KO mice might also demonstrate changes in pathways that regulate skeletal muscle energy metabolism. A well-described regulator of skeletal muscle energy metabolism is AMP activated protein kinase (AMPK), which alters both glucose and lipid metabolism in the setting of increased energy demand. We performed Western blots in the TA muscles from both chow and HFD fed WT and KO mice, which demonstrated that SOD2 KO muscles had an increased phosphorylation of the AMPKalpha subunit at the Thr172 activation site, compared to WT mice (**Figure 5A&5B**). Basal phosphorylation of AMPK was blunted in WT mice fed a HFD, however the increase in AMPK phosphorylation that was observed in KO mice fed a chow diet was abolished in KO mice fed a HFD. This result suggested that loss of SOD2 in a chow setting impacted on metabolic pathways that resulted in an increased need for energy, perhaps due to loss of mitochondrial efficiency, however this effect was confounded by the HFD milieu. In an attempt to further investigate the impact on muscle function, we performed qPCR analysis in the TA, EDL and Soleus muscles. There were no statistically significant differences in the abundance of any metabolism related genes in the KO (mCre+TAM) group compared to the three control groups (**Figure 5B–5D**). Moreover, given previous studies have indicated that developmental loss of SOD2 in skeletal muscle can lead to alterations in muscle morphology and branching ^21^, we subsequently analyzed the abundance of genes that might suggest changes in these pathways. Our data demonstrated that there was a change in the abundance of *Mef2c*, *Myog* and *Myod*, which are transcriptional regulators of skeletal muscle myogenesis (**Figure 5B–5D)**. Specifically, there was a trend towards reduced *Mef2C* and *MyoD* in the TA muscle, and in *MyoD* in the EDL. These findings are consistent with those previously described, and potentially indicate that SOD2 null muscle has a mild impairment in muscle myogenesis.

**Figure 5:**
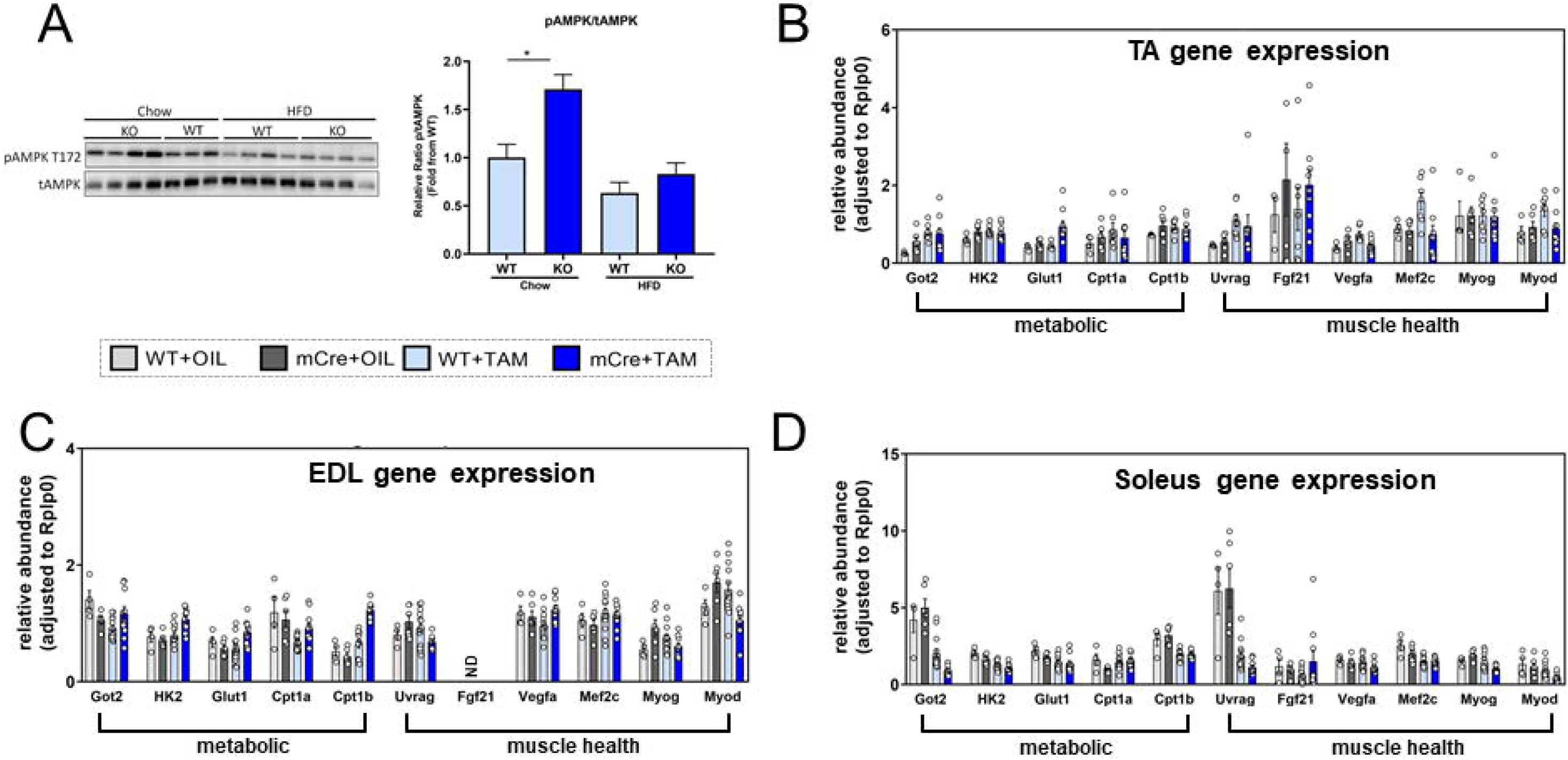
Analysis of AMP kinase activation and skeletal muscle gene expression in muscles from inducible, skeletal muscle specific SOD2 KO mice. (A) Western blot on protein from TA muscles isolated from WT+TAM (WT) and mCre+TAM (KO) mice fed a chow diet or HFD for phosphorylated AMP kinase (pAMPK T172), and total AMP kinase (tAMPK, loading control), and; (B) quantification of pAMPK/tAMPK abundance. qPCR analysis for genes involved in skeletal muscle metabolism and function from (C) TA (D) EDL and (E) soleus muscles from all four cohorts of HFD fed mice. Data presented as mean ± SEM. All data were compared by ANOVA with Fishers LSD post-hoc testing where * indicates a p-value<0.05. HFD = high fat diet, TA = *Tibialis anterior*, EDL = *Extensor digitorum longus*.

### Inducible, muscle specific knock out of SOD2 alters specific lipid pathways in skeletal muscle tissue

Finally, given that we have shown that inducible SOD2 deletion in skeletal muscle leads to subtle changes in mitochondrial and metabolic pathways, we sought to investigate if these changes affected lipid metabolism. Lipids are important signaling molecules, critical substrates for energy metabolism, and potent regulators of mitochondrial function. Thus, we performed ESI-MS/MS lipidomics analysis on muscle homogenates from all four cohorts of mice fed a HFD, and compared their lipidomes. We first analysed the total abundance of each of the 33 classes of lipids, which provided a global insight into the skeletal muscle lipidomes across the cohorts (**Figure 6A**). It was observed that in general the tamoxifen treatment had a noticeable impact on muscle lipid abundance, with an evident reduction in the abundance of several classes in mice treated with tamoxifen. This was particularly apparent in the triglyceride class (TG-SIM and TG-O), where these lipids appeared to be reduced by up to 50% as a response to Tamoxifen (WT+TAM and mCre-TAM cohorts; **Figure 6A**). Whilst many of the lipid classes were unaffected by the tamoxifen and deletion of SOD2, there were three particular lipid classes that were significantly reduced by SOD2 deletion, that were independent of any effects of Tamoxifen itself. These were diacylglycerols (DG), free fatty acids (FFA) and lysophospholipids (lysophosphatidylcholines - LPC, lysophosphatidylethanolamines - LPE and lysophosphatidylinositols - LPI). Several phosphate-containing lipid classes also exhibited a trend towards an increase in abundance in the SOD2 KO muscle, including phosphatidic acids (PA), phosphatidylcholines (PCs) and phosphatidylethanolamines (PEs). Given the significant differences observed in the DGs and FFAs, we performed a more in depth analysis of these lipids to investigate the effect of SOD2 KO on the individual species within these classes. We observed that several individual species of DG were significantly reduced by SOD2 KO, including many of the high abundance DGs such as those containing 16:χ and 18:χ fatty acids (**Figure 6B**). This was also reflected in the individual species of FFAs, where significant reductions were also observed for 4 different species including the 16:χ and 18:χ fatty acids (**Figure 6C**). The combined change in lysophospholipids demonstrated the global impact that SOD2 KO had on these lipid classes (**Figure 6D**). Overall, these data demonstrated that SOD2 deletion in skeletal muscle impacts on specific lipid pathways that are an important energy source for the mitochondria (i.e. FFAs). Moreover, there appeared to be a global impact on the level of phospholipids, with a potential activation of the pathways that clear the more toxic lysophospholipids such as LPC, LPE and LPI, into their acylated stable forms including PC and PE, a process catalyzed by lysophosphatidylcholine acyltransferase (LPCAT). In addition, an intriguing observation from these datasets is that the pathway responsible for conversion of phosphatidic acid (PA) into diacylglycerol (DG), appeared to be substantially impacted. This was evidenced by an increase in PA (the precursor) abundance and a decrease in DG (the downstream product) abundance, a reaction that is catalyzed by the enzyme phosphatidic acid phosphatase (PAP)/Lipin1. In light of these findings, we were interested to investigate whether we could detect differences in the amount Lipin1 in skeletal muscle of these animals. Western blotting for Lipin1 in WT and SOD2 KO muscles demonstrated that there were no alterations in the overall abundance of Lipin1 between WT and KO muscles, in either chow or HFD fed mice (**Figure 6E)**. Collectively, these findings suggest that SOD2 either directly or indirectly affects pathways that are involved in the metabolism of lipids, particularly phosphatidylcholine, providing interesting insights into a previously unexplored role of SOD2 in skeletal muscle.

**Figure 6:**
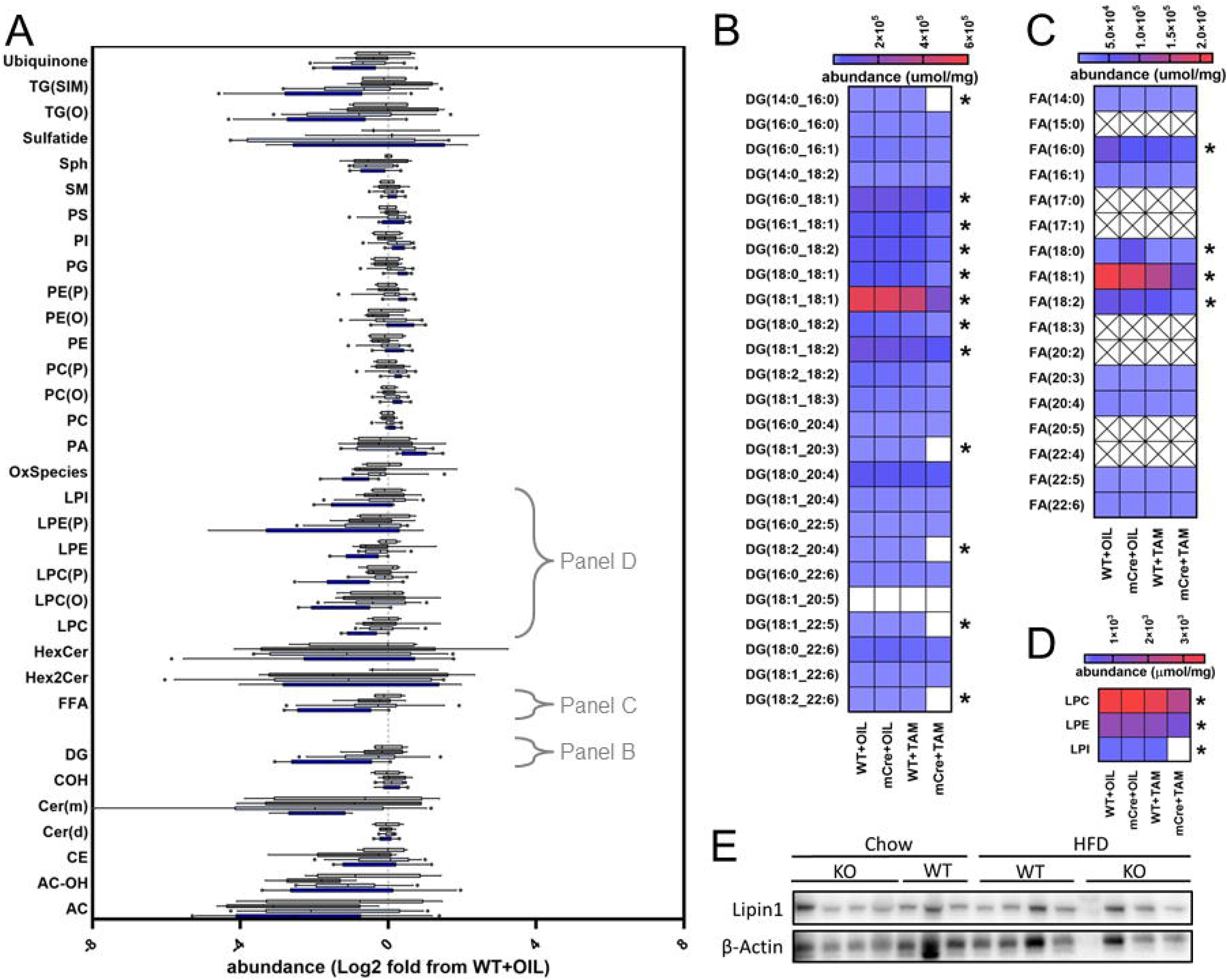
Analysis of lipid abundance in muscles from inducible, skeletal muscle specific SOD2 KO mice. Lipidomic assessment of lipid abundance in TA muscles from all four cohorts of HFD fed mice. (A) Total abundance of the 33 classes of lipid quantified in TA muscles across the four genotypes. Data are presented as log_2_ fold change from WT+TAM samples using box and whisker plots with black line representing the median, boxes representing the 25^th^ and 75^th^ percentile, whiskers represent 10-90^th^ percentile whilst individual dots represent values outside the 10^th^ and 90^th^ percentile. Heatmaps depicting abundance (μmol/mg) of individual species of (B) diacylglycerol (DG), (C) fatty acids (FA) and (D) total abundance of lysophosphatidylcholine (LPC), lysophosphatidylethanolamine (LPE) and lysophosphatidylinositol (LPI) lipids in TA muscle. Crossed boxes indicate species that were below the limit of detection. White boxes indicate species that fall below the lowest unit in the scale. Scale represents red as the highest abundance and light blue as the lowest abundance within a given range (shown on individual scale bars). (E) Western blot on protein from TA muscles isolated from WT+TAM (WT) and mCre+TAM (KO) mice fed a chow or high fat diet for Lipin1 and β-actin. Data in A-D were analysed by ANOVA with Benjamini-Hochberg correction (applied per sub-analysis) where * indicates a p-value<0.05.

## Discussion

In the current study, we demonstrate that post-developmental, skeletal muscle specific deletion of SOD2 imparts subtle effects on whole body physiology, which likely manifest as escalating deficits in mitochondrial function and subsequently energy metabolism. We also identify previously unrecognised alterations in specific lipid pathways in SOD2 KO muscles, which potentially define a new area of interest in the SOD2 field.

Previous studies investigating SOD2 deletion in skeletal muscle have done so either in heterozygous global null models^12^, or in muscle specific KO models that delete SOD2 whilst the muscles are still developing in utero^14,15^. Our study is the first to our knowledge that describes a deletion of SOD2 in skeletal muscle of mice that were already fully developed (i.e. at 8 weeks of age). Subsequent deletion of SOD2 for approximately 14 weeks (almost 60% of their entire life), did not recapitulate much of the prior phenotypes related to SOD2 deficiency in skeletal muscle, identifying important caveats to previous literature. The failure for our post-developmental model to recapitulate the robust muscle atrophy/weakness phenotype and impairments in glucose metabolism described in previous studies, speaks to the potential important roles of SOD2 in the developing muscle. Even when challenged with a high fat-diet (HFD), which we demonstrated to drive increases in oxidative damage and glucose intolerance, SOD2 KO mice did not demonstrate any additional deterioration.

Instead, the SOD2 KO mice demonstrated a more subtle phenotype, which, upon further investigation revealed a hereto unrecognised link between SOD2 and lipid metabolism. This included altered phospholipid (PL) abundance and the availability of important lipid intermediates such as diacylglyerols (DG) and free fatty acids (FFA). It is plausible that these pathways may have also been altered in previous models of SOD2 deletion, however it would have been difficult to tease out these subtle effects from the well progressed and overwhelming atrophy phenotype in those developmental models.

The alteration in these specific lipid species is intriguing, given that they all form part of a larger signaling network that regulates metabolism and several other important systems in skeletal muscle. Central to this network is the reciprocal regulation of phosphatidic acid (PA) and DG abundance, which is facilitated by opposing actions of PAP/lipin1 (PA → DG) and diacylglycerol kinase (DGK) (DG → PA)^22^. Several studies have investigated these specific enzyme complexes in skeletal muscle, all of which have demonstrated consistent phenotypes with that described in our and other muscle-specific SOD2 deletion studies. The data presented in our current study, would suggest that loss of SOD2 in skeletal muscle leads to either a reduction in PAP/Lipin1 activity, an increase in DGK activity, or a combination of both. Indeed, Lipin1 deficiency in skeletal muscle, which increases PA abundance and decreases DG abundance, leads to muscle atrophy, impaired autophagy and reductions in mitochondrial function^23–25^. This is consistent with the lipid alterations and reductions in mitochondrial protein abundance observed in SOD2 KO muscles. However, our data demonstrate that Lipin1 levels were not altered by SOD2 KO, which may indicate a lack of effect of this pathway. A caveat to these findings is that total abundance of Lipin1 is not a robust readout of activity, and thus other measures of function such as phosphorylation would strengthen these interpretations.

With regards to DGK, loss of different isoforms in skeletal muscle appears to impact on insulin signaling and AMP kinase activity, but not on mitochondrial function. Specifically, deletion of DGKδ increases DG levels, which in turn reduces AMPK phosphorylation and impaired lipid oxidation^26^. In SOD2 KO muscles, we observed decreased DG abundance and increased AMPK phosphorylation (in chow fed mice), consistent with effects described for DGK pathways. Unfortunately, we do not have data on DGK activity, so we cannot comment as to whether these pathways are responsible. Nevertheless, we speculate that in developed skeletal muscle, deletion of SOD2 initially impacts on this PA/DG lipid axis, which subsequently leads to reductions in mitochondrial function and a worsening lethargy phenotype, further perpetuating the effect of SOD2 deletion.

Further support for this PA/DG pathway being instrumental to the SOD2 phenotype, comes from studies performed in drosophila and *C. elegans*. Lin et al. described alterations in the DG/PA pathway that impacted on longevity via the ability of PA to activate the mTOR pathway^27^. These studies also demonstrated that modulation of these pathways resulted in higher susceptibility to oxidative stress-induced alterations in lifespan. Thus, PA/DG and oxidative stress may combine to form a critical axis that regulates skeletal muscle health and lifespan.

Despite these several lines of evidence, it remains unknown how the loss of SOD2 impacts on this axis, and perhaps disruption of redox signaling in skeletal muscle could influence PA levels and enzymes important in these pathways. Recent studies from Neufer et al. (2020) have demonstrated this to be a possibility, with studies from their lab linking beta-oxidation and redox homeostasis with Lipin1 activity and insulin resistance^28,29^. It is also possible that these proteins somehow interact with each other or within similar complexes, which might alter their activity or substrate availability. However, preliminary investigations from our group using deposited open-access protein-protein interaction datasets do not support this hypothesis (not shown), with no evidence for a direct or secondary interaction between these proteins. Another possibility is that loss of SOD2 subtly alters respiratory kinetics locally at the mitochondrial membrane, which secondarily impacts substrate and lipidome abundance within the mitochondria, leading to noticeable effects on cellular function over time. A recent study demonstrated that skeletal muscle specific loss of the enzyme Phosphatidylserine Decarboxylase (PSD), which synthesizes mitochondrial phosphatidylethaloamines (PE), leads to striking defects in mitochondrial and muscle function^30^. This loss of PE resulted in increased production of ROS from the mitochondria, implicating ROS as being important to this phenotype. Although we do not observe alterations in PE levels in this model, these data provide a direct role for mitochondrial lipids in regulating respiratory efficiency.

Overall, our data provide previously unrecognized effects of SOD2 deletion in skeletal muscle. Whilst our whole body analyses demonstrate only moderate impacts of post developmental deletion of SOD2 on some aspects of metabolism and muscle function compared to previous literature, our lipidomics analysis revealed intriguing alterations to the PA/DG pathway - providing a new link between redox biology and mitochondrial function. An obvious limitation to our current study is the short time frame over which we studied the animals, and had we allowed the animals to age for a longer period, a more robust phenotype may have developed. Moreover, we only challenged the animals with a HFD and no other interventions such as intense exercise training or muscle strain. Indeed, the HFD may not have been sufficient to precipitate the appropriate stress to accelerate disease. Conversely, had we utilized a more accelerated disease model, we may not have observed some of the subtle effects on lipid metabolism. Nevertheless, our current data provide unique insights into the underlying mechanisms by which SOD2 functions in skeletal muscle, highlighting previously unexplored interactions with specific lipid pathways important in development and disease.

## Methods

### Animals

All animal experiments were approved by the Alfred Research Alliance (ARA) Animal Ethics committee (E/1618/2016/B) and performed in accordance with the research guidelines set out by the National Health and Medical Research Council of Australia. SOD2 deletion was achieved using the Cre-Lox system. For inducible, skeletal muscle specific ablation, SOD2 floxed mice (C57BL/6J, a kind gift from Prof Takahiko Shimizu, Chiba University, Japan) were crossed with ACTA1-creERT2 mice (C57BL/6J background, Jackson Laboratories) to generate male cohorts of SOD2*^fl/fl^*-ACTA1-creERT2^+/-^ (SOD2 mCre) or SOD2*^fl/fl^*-ACTA1-creERT2^-/-^ (SOD2 WT). All mice were bred and sourced through the ARA Precinct Animal Centre and randomly allocated to groups. Cohorts of SOD2 mCre and WT mice were aged to 6-8 weeks of age before receiving oral gavage for 3 consecutive days of either Tamoxifen (80mg/kg) in sunflower oil, or sunflower oil alone. Following gavage, mice were left to recover for 2 weeks before being placed on high fat diet (43% energy from fat, #SF04-001 Specialty Feeds) for 12 weeks, or remained on normal chow diet (normal rodent chow, Specialty Feeds, Australia). Animals were housed at 22°C on a 12hr light/dark cycle with access to food and water *ad libitum* with cages changed weekly. At the end of the study mice were fasted for 4-6 hours and then anesthetized with a lethal dose of ketamine/xylazine before blood and tissues were collected, weighed and frozen for subsequent analysis.

### Tissue Sections and Nitrotyrosine Immunohistochemistry

TA muscles were carefully dissected and cut cross sectionally through the widest part of the muscle tissue. One half of the TA was embedded cut side down in OCT before being frozen in a bath of isopentane submerged in liquid nitrogen vapour. After freezing, blocks were brought to −20°C and 5μm sections were cut using a Leica Cryostat and then subjected to immunohistochemical staining for nitrotyrosine as described previously^31^. Briefly, mounted sections of TA muscle were fixed with cold acetone, and endogenous peroxidases were inactivated with 3% H_2_O_2_ in Tris-buffered saline. Sections were pre-blocked with a biotin-avidin blocking kit (Vector Laboratories) and then incubated with the nitrotyrosine antibody (Merck Milipore; 1:200) overnight at 4°C. Subsequent secondary antibody, biotinylated antirabbit immunoglobulin 1:100 (Dako) was added for 30 min, followed by horseradish peroxidase–conjugated streptavidin, diluted 1:500 (Dako), and incubated for 30 min in 3,3′-diaminobenzidine tetrahydrochloride (DAB) (Sigma-Aldrich). Images were captured on an Olympus Slide scanner VS120 (Olympus) and viewed in OlyVIA (Olympus, build 13771) and quantitated using a singular threshold setting in Fiji across all samples^32^.

### Peroxide Abundance using Amplex Red Assay

Muscle peroxide abundance was determined using the Amplex Red assay, as previously described^33^. Briefly, TA muscle tissue was homogenised using 20mM HEPES buffer (containing 1mM EGTA, 210mM Mannitiol and 70mM sucrose). After normalising protein concentration using a BCA assay (Pierce BCA assay kit), the quantification of hydrogen peroxide was determined using the Amplex Red Hydrogen Peroxide/Peroxidase Assay Kit (Molecular Probes) as per the manufacturer’s instructions.

### Glucose Tolerance Tests

Oral glucose tolerance tests (oGTT) were performed at different time points (0, 4 and 12 weeks post-HFD) throughout the study period at a dose of 1.5g/kg lean mass as determined by EchoMRI. All oGTTs were performed after a 5 hour fast as previously described^34^.

### EchoMRI

Body composition was analysed using the 4-in1 NMR Body Composition Analyzer for Live Small Animals, according to the recommendations of the manufacturer (EchoMRI LLC, Houston, TX, USA). This provides measurements of lean mass, fat mass and free water in living animals as previously described^34^.

### Whole Body Energetics

Mice were placed in the Promethion High-Definition Behavioral Phenotyping System for Mice (Sable Systems International, North Las Vegas, NV, USA) at 2 weeks post-tamoxifen (baseline) and 13 weeks post-Tamoxifen (11 weeks post HFD) of age for 3 consecutive days. Recordings for respirometry including energy expenditure (EE) and respiratory exchange ratio (RER) were collected over the final 24-hour period.

### Movement and Probability Mapping using Promethion

Behavioural phenotyping was conducted using assessments of activity monitoring (X, Y, Z beam breaks and wheel revolutions), in combination with food and water intake. The Promethion EthoScan utility created time and locomotion budgets with behavioural transition matrices for advanced behaviour analysis. Markov chain behaviour transition probability matrices were visualised utilising agl (Automatic Graph Layout, Microsoft Research; https://rise4fun.com/Agl/). Positional probability maps were generated across as an average of all positional locations during the 24-hour data collection period. Data was analysed and visualised in R (v3.5.3) with custom R scripts using the open-source SableBase package (Thomas Forester, 2016, Sable Systems International, Las Vegas, version 1.0).

### SDS-PAGE and Immunoblot

Skeletal muscle was lysed in radio-immunoprecipitation assay (RIPA) buffer supplemented with protease and phosphatase inhibitors. Matched protein quantities were separated by SDSPAGE and transferred to PVDF membranes. Membranes were blocked in 3% skim milk for 2 hours and then incubated with primary antibody overnight at 4°C for the following proteins: 4HNE (Abcam, ab46545), β-actin (Santa Cruz Biotech), Total OXPHOS Rodent WB Antibody Cocktail (MitoSciences), pan 14-3-3 (Santa Cruz), phospho-T172 AMPKalpha (Cell Signaling), Lipin-1 (Santa-Cruz Biotech) and total AMPKalpha (Cell Signaling). After incubation with primary antibodies, membranes were washed and probed with their respective HRP-conjugate secondary anti mouse or anti rabbit (Biorad) antibodies in 3% skim milk for 2 hours at room temperature, then visualised with enhanced chemiluminescent substrate (Pierce). Approximated molecular weights of proteins were determined from a co-resolved molecular weight standard (BioRad, #1610374). Image Lab Software (Bio-Rad) was used to perform densitometry analyses, and all quantification results were normalised to their respective loading control or total protein.

### Quantitative PCR (qPCR)

RNA was isolated from TA, EDL and Soleus muscles using RNAzol reagent and isopropanol precipitation. cDNA was generated from RNA using MMLV reverse transcriptase (Invitrogen) according to the manufacturer’s instructions. qPCR was performed on 10ng of cDNA using the SYBR-green method on a QuantStudio 7 Flex Real-Time PCR System, using primer sets outlined in Table 1. Primers were designed to span exon-exon junctions where possible, and were tested for specificity using BLAST (Basic Local Alignment Search Tool; National Centre for Biotechnology Information). Amplification of a single amplicon was estimated from melt curve analysis, ensuring only a single peak and an expected temperature dissociation profile were observed. Quantification of a given gene was determined by the relative mRNA level compared with control using the delta-CT method, which was calculated after normalisation to the housekeeping gene *Ppia* or *Rplp0*.

### Mitochondrial (mt)DNA to nuclear (n)DNA ratio

TA muscle tissue was homogenised in digestion buffer (100mM NaCl, 10mM Tris-HCl, 25mM EDTA, 0.5% SDS, pH 8.0) and then incubated in Proteinase K (250U/mL) for 1 hour at 55°C. Following this, total DNA was isolated using the phenol-chloroform extraction method. A qPCR reaction was then performed on 5ng of total DNA using a primer set that amplifies the mitochondrial gene mtCO3, and the genomic gene SDHA (see table 1 for primer sequences). Estimated abundance of each gene was used to generate a ratio of mitochondrial to nuclear DNA (mtDNA/nDNA), and this ratio was compared between genotypes.

### Lipidomics

Lipidomics was performed on approximately 50μg of soluble protein (homogenised and sonicated) from TA muscles taken from SOD2 WT+OIL, mCre+OIL, WT+TAM and mCre+TAM using LC electrospray ionisation MS/MS (LC-ESI-MS/MS) on an Agilent 6490 triple quadrupole (QQQ) mass spectrometer coupled with an Agilent 1290 series HPLC system and a ZORBAX eclipse plus C18 column as previously described^35^.

### Statistical Analyses

All data were expressed as mean ± standard error of the mean (SEM), except where otherwise stated (i.e. Figure 6A). Statistical comparisons in animal studies were analyzed by repeated measures 2-way ANOVA, two-way ANOVA with post-hoc testing, or one-way ANOVA with po-hoc testing as indicated in figure legends. Lipidomics, tissue analysis and cell based experiments were analyzed by either ANOVA with post-hoc testing (Fishers LSD) where appropriate, or paired students’ t-test unless otherwise stated. Analyses were performed using PRISM8 software and a p-value of p<0.05 was considered statistically significant.

### Data Inclusion and Exclusion Criteria

For animal experiments, phenotyping data points were excluded using pre-determined criteria if the animal was unwell at the time of analysis, there were technical issues identified (such as failed data acquisition in Promethion), values were biological implausible (such as RER=2.0) or data points that were identified as outliers using Tukey’s Outlier Detection Method (1.5IQR below Q1 or 1.5IQR above Q3). If repeated data points from the same mouse failed QC based on pre-determined criteria, or several data points were outliers as per Tukey’s rule, the entire animal was excluded from that given analysis (i.e. during glucose tolerance tests, indicating inaccurate dosing with gavage). For in vivo and in vitro tissue and molecular analysis, data points were only excluded if there was a technical failure (i.e. poor RNA quality, failed amplification in qPCR, failed injection in mass spectrometer), or the value was biological improbable. This was performed in a blinded fashion (i.e. on grouped datasets before genotypes were known).

## Acknowledgements

We acknowledge funding support from the Victorian State Government OIS program to Baker Heart & Diabetes Institute. BGD and ACC received support from the National Heart Foundation of Australia, Future Leader Fellowship scheme (101789 and 100067, respectively). We thank Prof Takahiko Shimizu from Chiba University, Japan for providing the floxed-mnSOD mice. We also thank all members of the MMA, LMCD and Metabolomics laboratories at BHDI for their ongoing contributions.

## Author contributions

BGD conceived and designed the study, and wrote the manuscript. BGD, AZ, CY, YL, YT, STB, SW, TS performed all experiments. AS and JBdeH performed and analysed Amplex Red and Nitrotyrosine assays. PJM provided expertise in lipidomics analysis. MTC provided access to floxed mice and ongoing research support. ACC provided reagents, experimental advice and access to resources. All authors read and approved the manuscript.

## References

1. Wanagat, J. & Hevener, A.L. Mitochondrial quality control in insulin resistance and diabetes. Curr Opin Genet Dev 38, 118–126 (2016).

2. Montgomery, M.K. & Turner, N. Mitochondrial dysfunction and insulin resistance: an update. Endocrine connections 4, R1–R15 (2015).

3. Trifunovic, A., et al. Premature ageing in mice expressing defective mitochondrial DNA polymerase. Nature 429, 417–423 (2004).

4. Giacco, F. & Brownlee, M. Oxidative stress and diabetic complications. Circ Res 107, 1058–1070 (2010).

5. Brownlee, M. The pathobiology of diabetic complications: a unifying mechanism. Diabetes 54, 1615–1625 (2005).

6. Kauppila, T.E.S., Kauppila, J.H.K. & Larsson, N.G. Mammalian Mitochondria and Aging: An Update. Cell metabolism 25, 57–71 (2017).

7. Andreux, P.A., Houtkooper, R.H. & Auwerx, J. Pharmacological approaches to restore mitochondrial function. Nature reviews. Drug discovery 12, 465–483 (2013).

8. Boden, M.J., et al. Overexpression of manganese superoxide dismutase ameliorates high-fat diet-induced insulin resistance in rat skeletal muscle. American journal of physiology. Endocrinology and metabolism 303, E798–805 (2012).

9. Hoehn, K.L., et al. Insulin resistance is a cellular antioxidant defense mechanism. Proceedings of the National Academy of Sciences of the United States of America 106, 17787–17792 (2009).

10. Li, Y., et al. Dilated cardiomyopathy and neonatal lethality in mutant mice lacking manganese superoxide dismutase. Nature genetics 11, 376–381 (1995).

11. Crane, J.D., et al. Elevated mitochondrial oxidative stress impairs metabolic adaptations to exercise in skeletal muscle. PloS one 8, e81879 (2013).

12. Kang, L., et al. Heterozygous SOD2 deletion impairs glucose-stimulated insulin secretion, but not insulin action, in high-fat-fed mice. Diabetes 63, 3699–3710 (2014).

13. Koyama, H., et al. Antioxidants improve the phenotypes of dilated cardiomyopathy and muscle fatigue in mitochondrial superoxide dismutase-deficient mice. Molecules 18, 1383–1393 (2013).

14. Kuwahara, H., et al. Oxidative stress in skeletal muscle causes severe disturbance of exercise activity without muscle atrophy. Free radical biology & medicine 48, 1252–1262 (2010).

15. Lustgarten, M.S., et al. Conditional knockout of Mn-SOD targeted to type IIB skeletal muscle fibers increases oxidative stress and is sufficient to alter aerobic exercise capacity. American journal of physiology. Cell physiology 297, C1520–1532 (2009).

16. Ikegami, T., et al. Model mice for tissue-specific deletion of the manganese superoxide dismutase (MnSOD) gene. Biochemical and biophysical research communications 296, 729–736 (2002).

17. Izuo, N., et al. Brain-Specific Superoxide Dismutase 2 Deficiency Causes Perinatal Death with Spongiform Encephalopathy in Mice. Oxidative medicine and cellular longevity 2015, 238914 (2015).

18. Han, Y.H., et al. Adipocyte-Specific Deletion of Manganese Superoxide Dismutase Protects From Diet-Induced Obesity Through Increased Mitochondrial Uncoupling and Biogenesis. Diabetes 65, 2639–2651 (2016).

19. Kawakami, S., et al. Antioxidant, EUK-8, prevents murine dilated cardiomyopathy. Circulation journal : official journal of the Japanese Circulation Society 73, 2125–2134 (2009).

20. Liu, G., et al. Bladder function in mice with inducible smooth muscle-specific deletion of the manganese superoxide dismutase gene. American journal of physiology. Cell physiology 309, C169–178 (2015).

21. Ahn, B., et al. Mitochondrial oxidative stress impairs contractile function but paradoxically increases muscle mass via fibre branching. Journal of cachexia, sarcopenia and muscle 10, 411–428 (2019).

22. Brindley, D.N., Pilquil, C., Sariahmetoglu, M. & Reue, K. Phosphatidate degradation: phosphatidate phosphatases (lipins) and lipid phosphate phosphatases. Biochimica et biophysica acta 1791, 956–961 (2009).

23. Zhang, P., Verity, M.A. & Reue, K. Lipin-1 regulates autophagy clearance and intersects with statin drug effects in skeletal muscle. Cell metabolism 20, 267–279 (2014).

24. Schweitzer, G.G., et al. Loss of lipin 1-mediated phosphatidic acid phosphohydrolase activity in muscle leads to skeletal myopathy in mice. FASEB journal : official publication of the Federation of American Societies for Experimental Biology 33, 652–667 (2019).

25. Rashid, T., et al. Lipin1 deficiency causes sarcoplasmic reticulum stress and chaperone-responsive myopathy. The EMBO journal 38(2019).

26. Jiang, L.Q., et al. Diacylglycerol kinase-delta regulates AMPK signaling, lipid metabolism, and skeletal muscle energetics. American journal of physiology. Endocrinology and metabolism 310, E51–60 (2016).

27. Lin, Y.H., et al. Diacylglycerol lipase regulates lifespan and oxidative stress response by inversely modulating TOR signaling in Drosophila and C. elegans. Aging cell 13, 755–764 (2014).

28. Smith, C.D., Schmidt, C.A., Lin, C.T., Fisher-Wellman, K.H. & Neufer, P.D. Flux through mitochondrial redox circuits linked to nicotinamide nucleotide transhydrogenase generates counterbalance changes in energy expenditure. The Journal of biological chemistry 295, 16207–16216 (2020).

29. Smith, C.D., et al. Genetically increasing flux through beta-oxidation in skeletal muscle increases mitochondrial reductive stress and glucose intolerance. American journal of physiology. Endocrinology and metabolism (2021).

30. Heden, T.D., et al. Mitochondrial PE potentiates respiratory enzymes to amplify skeletal muscle aerobic capacity. Science advances 5, eaax8352 (2019).

31. Sharma, A., et al. Direct Endothelial Nitric Oxide Synthase Activation Provides Atheroprotection in Diabetes-Accelerated Atherosclerosis. Diabetes 64, 3937–3950 (2015).

32. Schindelin, J., et al. Fiji: an open-source platform for biological-image analysis. Nat Methods 9, 676–682 (2012).

33. Sharma, A., et al. The nuclear factor (erythroid-derived 2)-like 2 (Nrf2) activator dh404 protects against diabetes-induced endothelial dysfunction. Cardiovasc Diabetol 16, 33 (2017).

34. Bond, S.T., Kim, J., Calkin, A.C. & Drew, B.G. The Antioxidant Moiety of MitoQ Imparts Minimal Metabolic Effects in Adipose Tissue of High Fat Fed Mice. Frontiers in physiology 10, 543 (2019).

35. Huynh, K., et al. High-Throughput Plasma Lipidomics: Detailed Mapping of the Associations with Cardiometabolic Risk Factors. Cell chemical biology 26, 71–84 e74 (2019).

